# *SELF-PRUNING* affects auxin responses synergistically with the cyclophilin A DIAGEOTROPICA in tomato

**DOI:** 10.1101/271387

**Authors:** Willian B. Silva, Mateus H. Vicente, Jessenia M. Robledo, Diego S. Reartes, Renata C. Ferrari, Ricardo Bianchetti, Wagner L. Araújo, Luciano Freschi, Lázaro E. P. Peres, Agustin Zsögön

## Abstract

**Summary:** The antiflorigenic signal SELF-PRUNING, which controls growth habit, exerts its effects through auxin transport, signaling and metabolism in tomato.

**Abstract:** The *SELF PRUNING* (*SP*) gene is a key regulator of growth habit in tomato (*Solanum lycopersicum*). It is an ortholog of *TERMINAL FLOWER 1*, a phosphatidyl-ethanolamine binding protein with anti-florigenic activity in *Arabidopsis thaliana*. A spontaneous loss-of-function *sp* mutation has been bred into a large number of industrial tomato cultivars, as it produces a suite of pleiotropic effects that are favorable for mechanical harvesting, including determinate growth habit, short plant stature and simultaneous fruit ripening. However, the physiological basis for these phenotypic differences has not been thoroughly explained. Here, we show that the *sp* mutation alters polar auxin transport as well as auxin responses such gravitropic curvature and elongation of excised hypocotyl segments. We further demonstrate that free auxin levels and auxin-regulated gene expression patterns are altered in *sp*, with epistatic effects of *diageotropica*, a mutation in a cyclophilin A protein-encoding gene. Our results indicate that SP impacts growth habit in tomato, at least in part, via changes in auxin transport and responsiveness. These findings hint at novel targets that could be manipulated in the control of growth habit and productivity.

## Introduction

Shoot architecture is a key agricultural trait determined mainly by side branching, internode elongation and shoot determinacy (Wang and Li, 2008). Each of these parameters configures an active research area where considerable theoretical and applied knowledge has been gained over the last decade. Shoot determinacy is a domestication trait in crop species as diverse as soybean (*Glycine max*), common bean (*Phaseolus vulgaris*) and tomato (*Solanum lycopersicum*) (Pnueli et al., 1998; Tian et al., 2010; Repinski et al., 2012). Tomato is a perennial species cultivated as annual. Wild tomatoes display indeterminate growth, resulting from a sequential addition of modules (sympodial units) formed by three leaves and an inflorescence. Sympodial growth starts in tomato when the vegetative apical meristem is converted into floral after a series of 8-12 internodes with leaves (Samach and Lotan, 2007). Vegetative growth, however, continues through the topmost axillary meristem, which grows vigorously displacing the inflorescence to the side and producing a new sympodial unit with three leaves and an inflorescence. This process is indefinitely iterated by concatenation of sympodial units one on top of the other. A spontaneous recessive mutant with compact, bushy growth habit and a reduced number of leaves in successive sympodial units was discovered in 1914 (Yeager, 1927; MacArthur, 1934). It was later shown that the mutation is a single nucleotide substitution in the *SELF-PRUNING* (*SP*) gene (Pnueli et al., 1998), which shares sequence similarity with a group of mammalian polypeptides involved in cell signaling, phosphatidylethanolamine binding proteins (PEBPs) (Hengst et al., 2001; Kroslak et al., 2001). Breeding of this mutation into industrial tomato cultivars was instrumental in the advent of mechanical harvest (Rick, 1978; Stevens and Rick, 1986). The loss-of-function *sp* mutant leads to determinate growth habit, as opposed to the indeterminate growth habit of wild-type tomatoes. The determinate growth habit occurs via progressively reduced number of leaves per sympodium until termination in two consecutive inflorescences that top vertical growth of the plant (Samach and Lotan, 2007). Hence, this phenotype leads to simultaneous fruit ripening, therefore allowing mechanical harvest in field-grown processing tomatoes (Stevens and Rick, 1986).

*SP* belongs to the *CETS* gene family, which comprises *CENTRORADIALIS* (*CEN*) and *TERMINAL FLOWER 1* (*TFL1*) of *Antirrhinum* and *Arabidopsis*, respectively (Wickland and Hanzawa, 2015). *SINGLE FLOWER TRUSS (SFT)/SP3D* - a homolog of *FLOWERING LOCUS T* (*FT*) and *HEADING DATE 3A* (*HD3A*) in Arabidopsis and rice, respectively - is another *CETS* gene involved in the control of growth habit in tomato (Alvarez et al., 1992; Kojima et al., 2002). Unlike *sp* mutants, which do not affect flowering time, tomato *sft* loss-of-function mutants are late flowering, and also show a disruption in sympodial growth pattern: they produce a single and highly vegetative inflorescence, alternating solitary flowers and leaves (Molinero-Rosales et al., 2004). The final phenotypic outcome produced by SP and SFT depends on their local ratio, with the former maintaining meristems in an indeterminate state and the latter promoting the transition to flowering (Park et al., 2014). Heterozygous *sft* mutants in a homozygous *sp* mutant background display yield heterosis in tomato (Krieger et al., 2010). Hence, the *SP/SFT* genetic module has been proposed as a target to increase crop yield via changes in plant architecture (McGarry and Ayre, 2012; Zsögön et al., 2017). It has also been previously suggested that *SP* function could be linked to auxin (Pnueli *et al.*, 2001), a hormone with strong effects on plant morphogenesis (Berleth and Sachs, 2001).

Auxin is a key controller of plant development; however, its role in the regulation of plant growth habit is still unclear. An aspect that sets auxin apart from other plant hormones is the relatively well understood nature of its transport through the plant body (Friml, 2003; Petrášek and Friml, 2009). Polar auxin transport (PAT), which occurs basipetally from the apical meristem, is critically important for the distribution of auxin within plant tissues (Rubery and Sheldrake, 1974; Sheldrake, 1974). PAT works as an organizer of apical-basal polarity in the plant body (Friml et al., 2006), thus controlling a multiplicity of developmental processes (Reinhardt et al., 2003; Blilou et al., 2005; Scarpella et al., 2006).

It has recently been shown that the cyclophilin A protein DIAGEOTROPICA (DGT) affects polar auxin transport (PAT) in tomato (Ivanchenko et al., 2015). DGT is a cyclophilin A protein with peptidyl-prolyl trans-cis isomerase (PPIase) enzymatic activity (Takahashi et al., 1989; Oh et al., 2006). Cyclophilins catalyse not only rate-limiting steps in the protein folding pathway but can also participate in the folding process as molecular chaperones (Kumari et al., 2013). DGT function is highly conserved across plant taxa (Lavy et al., 2012). In tomato, some of the most significant phenotypic defects caused by the lack of functional DGT protein are horizontal shoot growth, thin stems, altered secondary vascular differentiation and roots lacking lateral branches (Zobel, 1973; Muday et al., 1995; Coenen et al., 2003). Here, we investigated whether *SP* affects auxin responses, by itself, and in combination with *DGT*. We produced four combinations of functional and loss-of-function mutant alleles of *SP* and *DGT* (*i.e. SP DGT*, *SP dgt*, *sp DGT* and *sp dgt*) in a single tomato genetic background (cv. Micro-Tom) and assessed a series of physiological responses to auxin. We found that free auxin levels, polar auxin transport and gravitropic curvature of the shoot apex are all altered by *SP*. Our results further show that *SP* and *DGT* reciprocally affect *AUX/IAA* and *ARF* transcript abundances at the sympodial meristem, the key niche of *SP* function in growth habit.

## Results

Comparison of the four combinations of homozygous wild-type and mutant lines for *SP* and *DGT* (*i.e.*, *SP DGT*, *SP dgt*, *sp DGT* and *sp dgt*), showed that growth habit was affected solely by *SP* and not by *DGT* (Fig. 1). Regardless of their *DGT* or *dgt* allele, *SP* plants showed indeterminate growth whereas *sp* mutants were always determinate (Fig. 1). Time to flowering, however, was affected by both genes in combinatorial fashion. *dgt* plants flowered late, independently of the *SP* allele (Fig. 1). The *sp DGT* genotype showed consistently precocious flowering, and this was confirmed in an independent experiment by analysis of the rate of shoot apical meristem maturation (Fig S1). The number of leaves to the first inflorescence was also affected by the combination of alleles (Fig. 1), albeit not reflecting the time to flowering. The *dgt* mutant produced more leaves before flowering, but this effect was abolished in the double mutant *sp dgt*. Regardless of their *SP* allele, *dgt* mutants exhibited markedly reduced transcript abundance of the flowering inducer *SINGLE FLOWER TRUSS (SFT*) compared to *DGT* plants (Fig. 1), which fits with the delayed flowering in these mutants in both *SP* and *sp* backgrounds (Fig. 1).

**Figure 1.**
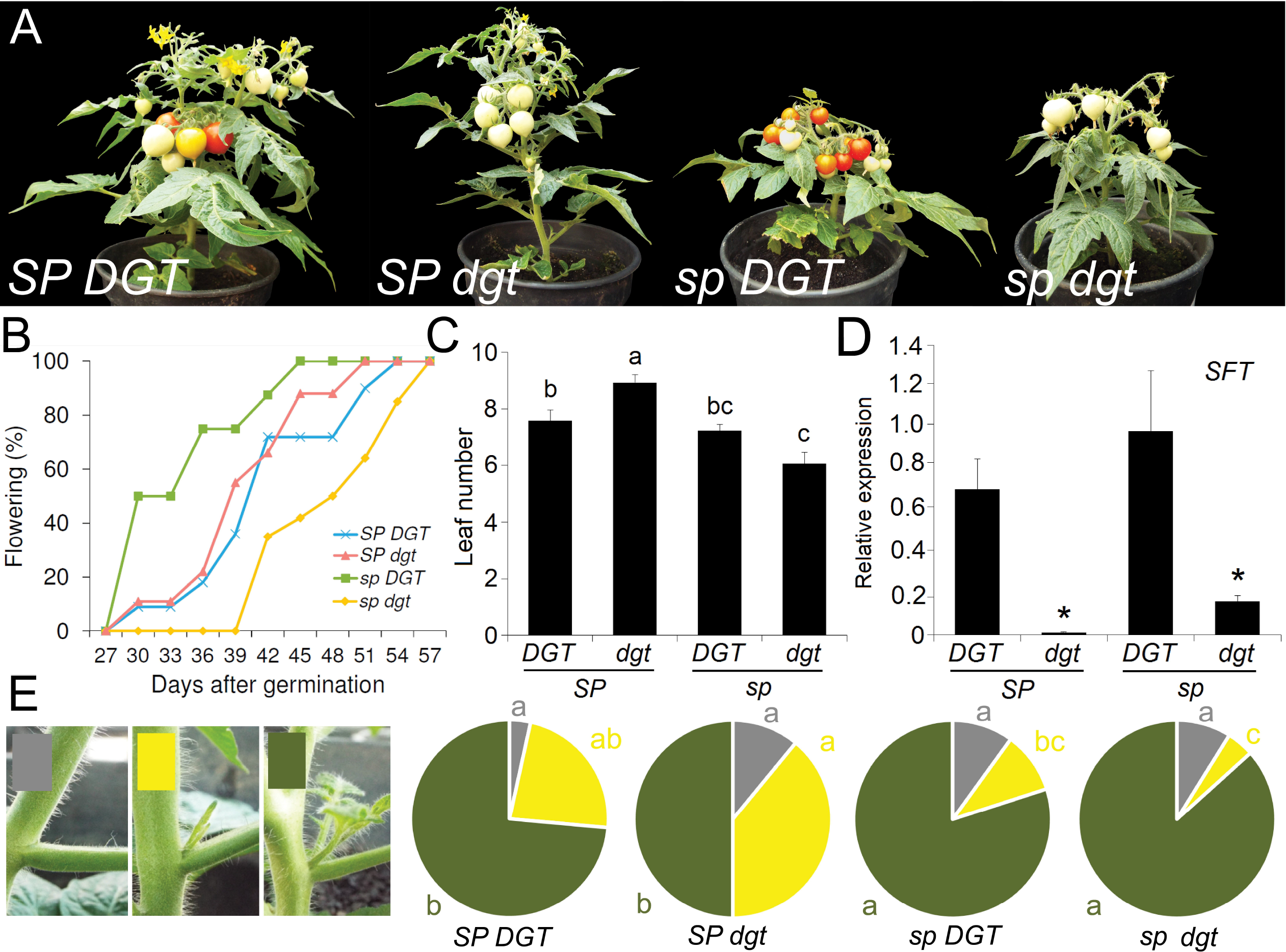
Additive phenotype of the *self-pruning* (*sp*) and *diageotropica* (*dgt*) mutations in tomato cv. Micro-Tom (MT). (A) Representative plants of *SP DGT*; *SP dgt*; *sp DGT* (cv. MT) and *sp dgt*, 90 dag. Note the simultaneous fruit ripening in *sp* compared to *SP*, a well-known effect of the *sp* mutation. The *dgt* mutation delays fruit ripening (at least in part due to its late flowering, as indicated in b) in either genetic background. Bar=10 cm. **(B) Chronological time to flowering in sp and *dgt* mutants.** Percentage of plants (*n*=15) with at least one open flower. MT (*sp DGT*) plants flower earlier than wild type (*SP DGT*), whereas dgt mutants are late flowering. **(C) Developmental time to flowering in *sp* and *dgt* mutants.** The number of leaves produced before the first inflorescence was reduced in *sp DGT* (MT) and increased in genotypes carrying the functional allele of *SP*. Letters indicate statistically significant differences (Dunn’s multiple comparisons test p<0.05). **(D)*sp* and *dgt* alter expression of the flowering inducer *SINGLE FLOWER TRUSS* (*SFT*).** The *dgt* mutation leads to lower *SFT* expression and thus delays flowering. A minor influence from *SP* reducing *SFT* levels is also noticeable. Asterisks indicate statistically significant differences with the wild-type *SP DGT* (Student’s t-test, p<0.05). **(E) Effect of *sp* and *dgt* on side branching.** Schematic representation of side branching in shoots of *SP DGT*; *SP dgt*; *sp DGT* (MT) and *sp dgt* (*n*=15). Pie charts depicting the distribution of side branches in each genotype 60 dag. Grey denotes absence of axillary bud; yellow, a visible bud (>1cm) and dark green, a full branch (with one or multiple leaves). Letters indicate statistically significant differences (Dunn’s multiple comparisons test p<0.05).

The tomato cultivar Micro-Tom (MT) harbors a mutation in *DWARF* (*D*), a gene coding for a key enzyme in the brassinosteroid biosynthesis pathway (Bishop et al., 1999). Since brassinosteroids are known to influence the flowering induction network (Domagalska et al., 2010; Li et al., 2010), we ascertained whether *D* could be influencing the effects of *SP* on flowering time. Using a near-isogenic MT line harboring the functional *D* allele (Carvalho et al., 2011), we constructed four allelic combinations of *SP* and *D* (*i.e. SP D*, *SP d*, *sp D* and *sp d*, Fig S2) and assessed their flowering time. The results show an effect of *D* on flowering time (Fig. S2) but not on the number of leaves produced to the first inflorescence, which was again reduced exclusively by the presence of the *sp* mutant allele (Fig. S2). Axillary branching was affected mainly by the *SP* gene, which led to reduced bud outgrowth in plants carrying the wild-type allele; the *dgt* mutation, however, exacerbated this repressing effect (Fig. 1). *sp* mutants, on the other hand, branched more profusely when combined with *dgt* than *DGT* (Fig. 1). Thus, *dgt* can enhance apical dominance or increase branching, depending on the presence or absence of a functional *SP* allele, respectively. The number of leaves on the primary shoot was increased by *SP*, regardless of the *DGT* allele (Table 1). Plant height was additively controlled by both genes, whereas no difference between genotypes was found in the length of the fourth internode or leaf insertion angle (Table 1). Stem diameter was increased by functional *DGT*, irrespective of the *SP* allele (Table 1). The number of inflorescences was synergistically determined by both genes, whereby pairing of functional *SP* and *DGT* led to an increased number compared to all other allele combinations (Table 1).

**Table 1.**
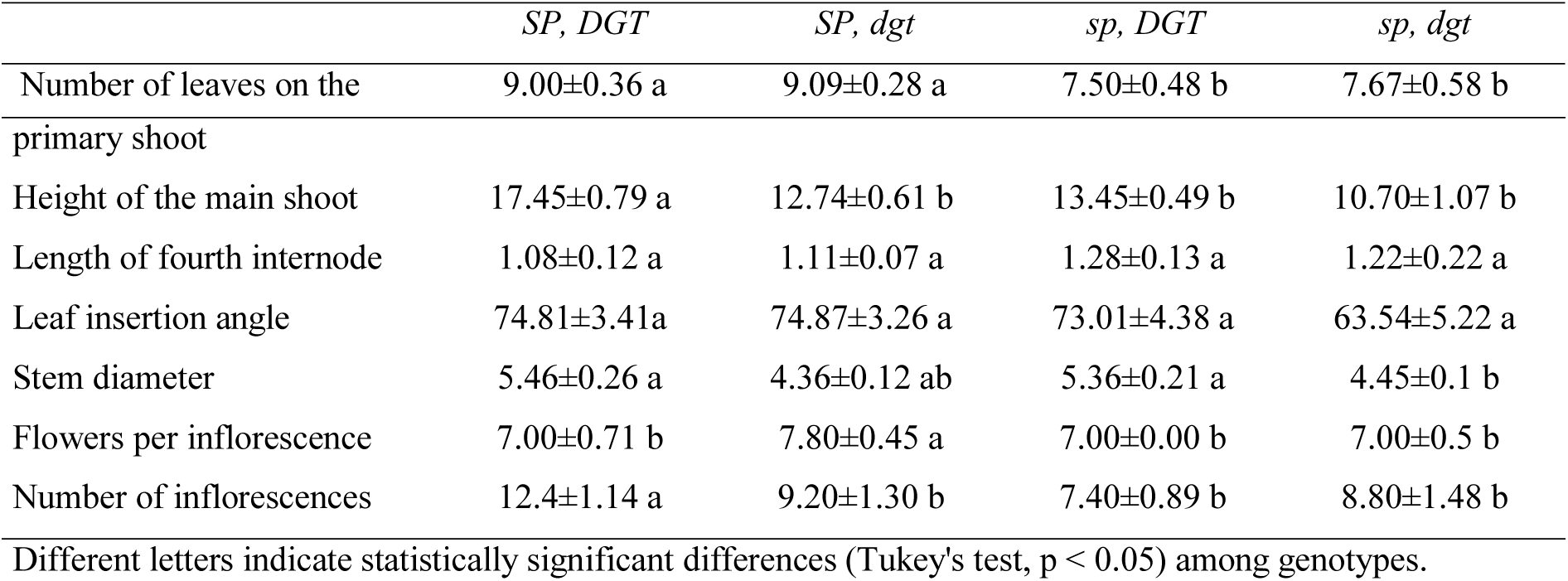
Parameters that define growth habit: (i) number of leaves on the primary shoot (PS) (i.e. number of leaves up to the first inflorescence); (ii) number of leaves on the main shoot (MS) (i.e. number of leaves of PS plus leaves on sympodial units (SU) following the first inflorescence); (iii) height of PS, MS and lateral shoot (LS); iv) internode length (cm); v) leaf angle insertion; vi) diameter of stem and vii) number of flowers, fruits and flowers per inflorescence. Measurements performed 60 days after germiantion. Data are mean ± s.e.m. (n = 10 plants).

|  | *SP, DGT* | *SP, dgt* | *sp, DGT* | *sp, dgt* |
| --- | --- | --- | --- | --- |
| Number of leaves on the | 9.00±0.36 a | 9.09±0.28 a | 7.50±0.48 b | 7.67±0.58 b |
| primary shoot |  |  |  |  |
| Height of the main shoot | 17.45±0.79 a | 12.74±0.61 b | 13.45±0.49 b | 10.70±1.07 b |
| Length of fourth internode | 1.08±0.12 a | 1.11±0.07 a | 1.28±0.13 a | 1.22±0.22 a |
| Leaf insertion angle | 74.81±3.41a | 74.87±3.26 a | 73.01±4.38 a | 63.54±5.22 a |
| Stem diameter | 5.46±0.26 a | 4.36±0.12 ab | 5.36±0.21 a | 4.45±0.1 b |
| Flowers per inflorescence | 7.00±0.71 b | 7.80±0.45 a | 7.00±0.00 b | 7.00±0.5 b |
| Number of inflorescences | 12.4±1.14 a | 9.20±1.30 b | 7.40±0.89 b | 8.80±1.48 b |

Next, the endogenous levels of free indolyl-3-acetic acid (IAA), which is the most abundant auxin in plants (Bartel and Fink, 1995), was determined in three sections of tomato seedlings: leaves plus cotyledons, hypocotyls and roots (Fig. 2). In leaves plus cotyledons, IAA concentration was more than twice higher in *sp dgt* double mutant than in the other three genotypes (Fig. 2). In the hypocotyl tissues, *SP DGT* seedlings had the lowest free IAA content, the *sp* and *dgt* single mutants presented intermediate values and the double mutant (*sp dgt*) exhibited the highest IAA levels (Fig. 2). Although root IAA levels were clearly higher than in the other hypocotyl regions analyzed, no statistically significant differences in root IAA content was observed between the four genotypes (Fig. 2).

**Figure 2.**
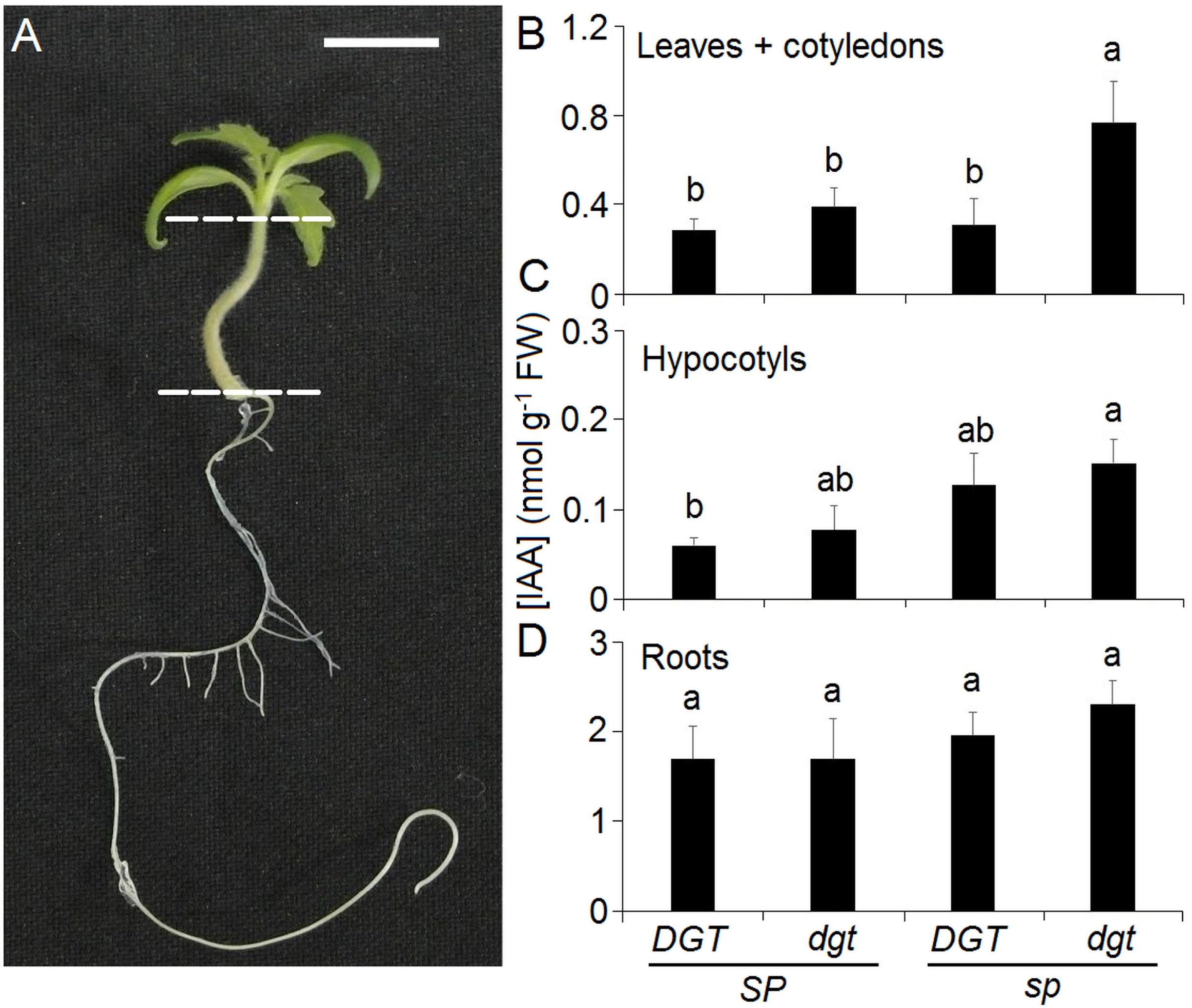
Auxin levels in tomato seedlings are affected synergistically by the *self-pruning* (*sp*) and *diageotropica (dgt)* mutations. (A) Representative 7-day old seedling showing the dissection points for auxin quantitation. Free IAA levels in (B) leaves + cotyledons, (C) hypocotyls and (D) roots. Data are mean±s.e.m. (*n*=10) Different letters indicate statistically significant differences (Tukey’s test, p<0.05) among genotypes.

To understand the variation in endogenous free IAA levels within the seedling tissues and among the four genotypes, we next quantified polar auxin transport (PAT) in hypocotyl segments. PAT was highest in *SP DGT*, intermediate in *sp* and *dgt* single mutants, and lowest in the double mutant (Fig. 3). This indicates that both *sp* and *dgt* alleles reduce PAT and that their effects were additive. As PAT and auxin concentration are known to influence vascular patterning (Scarpella, 2017), we also analyzed xylem anatomy in cross-sections of stems in adult plants (Fig 3). Quantification of xylem vessel density and mean vessel size revealed an antagonistic relationship between *SP* and *DGT*. Whereas *SP* tends to reduce vessel density and increase their size, *DGT* increases vessel density with concomitantly lower vessel sizes (Fig. 3). These results, however, obscure a more complex pattern, which is revealed when analyzing the vessel size distributions. The functional *DGT* allele increased the incidence of larger (>800 μm^2^ cross-sectional area) vessels, particularly in *sp* mutant plants (Fig 3). Another physiological response affected by PAT is negative gravitropism of the shoot (Morita, 2010). The kinetics of gravitropic curvature in seedling shoots was affected by both *SP* and *DGT* (Fig. 4). Loss of *SP* function decreased gravitropic response in both *DGT* and *dgt* backgrounds. Hypocotyl elongation in response to exogenous auxin and *in vitro* rhizogenesis from cotyledon explants are assays to determine auxin sensitivity (Cary et al., 2001). The *dgt* mutation considerably reduces hypocotyl responsivity to auxin in all concentrations, as described previously (Kelly and Bradford, 1986; Rice and Lomax, 2000). The functional *SP* allele increased hypocotyl elongation in a *DGT* background and also exerts a significant compensatory effect on the elongation response in the *dgt* mutant (Fig. 4). In the *in vitro* root regeneration assay, as expected, root formation was reduced in *dgt* mutants (Coenen and Lomax, 1998), but also in *sp* compared to *SP* in the presence of a functional *DGT* allele (Fig. S3).

**Figure 3.**
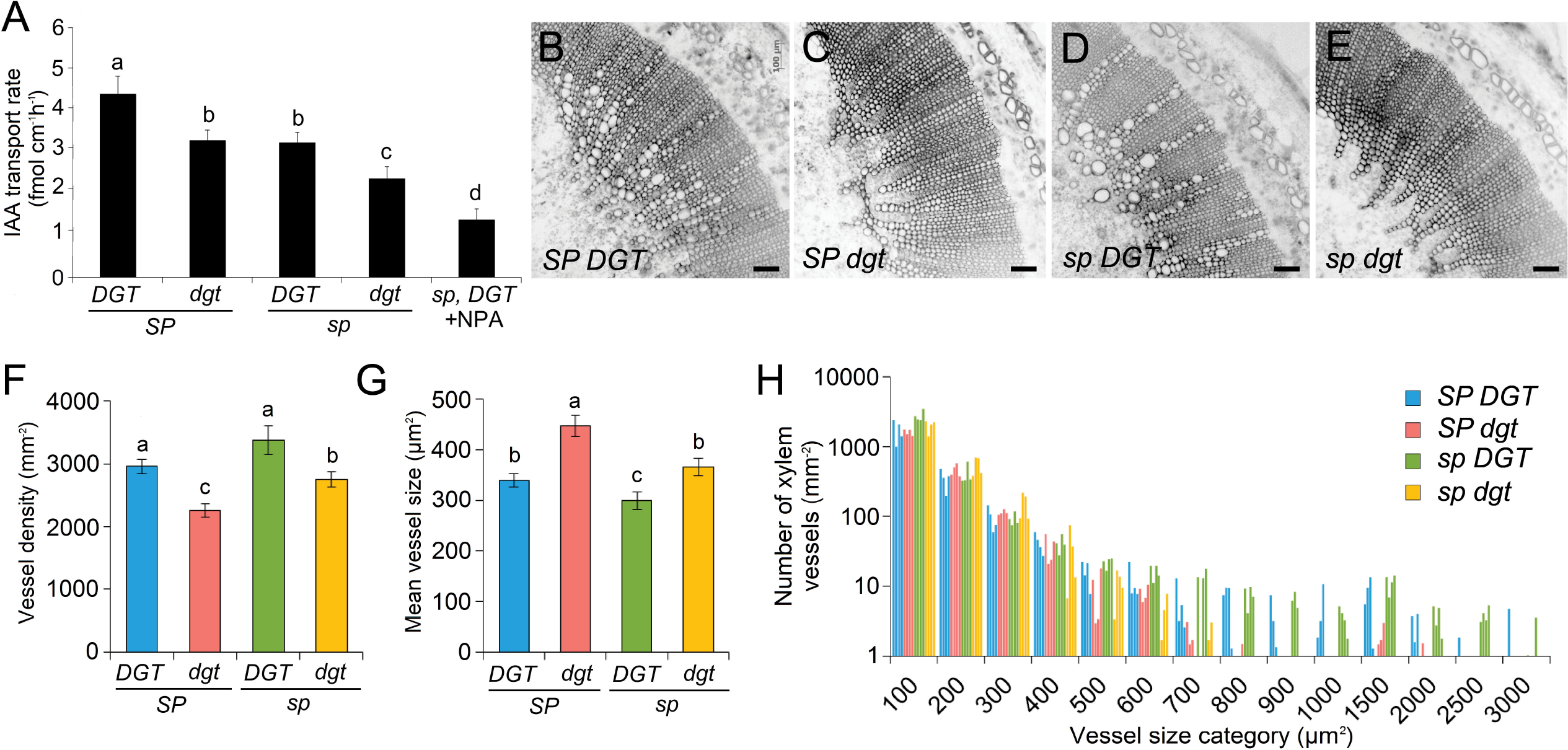
(A) The *self-pruning* (*sp*) mutation exacerbates defective polar auxin transport in hypocotyls caused by *diageotropica* (*dgt*). Basipetal ^3^H-IAA transport in 10 mm hypocotyl sections of wild-type (*SP*, *DGT*), *SP dgt*, *sp DGT* (cv. Micro-Tom; also the negative control treated with NPA) and double mutant *sp dgt* roots. Data are mean±s.e.m. (*n*=10). Asterisk indicates statistically significant differences between treatments (ns, non-significant; **p≤0.05; \*\*\**p*≤0.01, t-test). **(B-E) Vascular patterning in *sp* and *dgt* stems.** Crosssections of the fifth internode taken 45 dag. Bar = 100 μm. **(F) Vessel density and (G) mean vessel size in *sp* and *dgt* stems.** Letters indicate significant differences (p<0.05 ANOVA, Tukey). **(H) Vessel size distribution in the xylem of *sp* and *dgt* mutants.** The x-axis shows the upper values of cross-sectional area for each vessel size category. The bars within each category represent a single individual plant (n=4 per genotype).

Histochemical analysis of *DR5* promoter activity revealed no discernible staining difference in both *SP* and *sp* seedlings incubated in water, although roots of the *sp* mutant showed a shorter trace of GUS precipitate in the vascular cylinder (Fig. 5). Exogenous IAA, however, strongly induced GUS expression in *SP* compared to *sp* plants, which was evident both in seedlings and in root tips and confirmed by fluorimetric GUS quantitation (Fig. 5). Fainter GUS staining was observed for both auxin treated and untreated roots in the *dgt* mutant (Fig S4). As PIN-FORMED (PIN) auxin efflux transporters are key players determining auxin distribution in plants, we quantified the relative expression of the *PIN1, PIN2* and *PIN3* genes in roots with or without prior auxin incubation. Auxin treatment induced *PIN1* and *PIN3* expression in all genotypes, except in the *sp dgt* double mutant (Fig. 5). *PIN2* expression was reduced by auxin incubation in *SP DGT, SP dgt* and *sp dgt*, but not in *sp DGT* (*i.e*. cv Micro-Tom), where a low basal level of expression was observed for all three genes.

**Figure 4.**
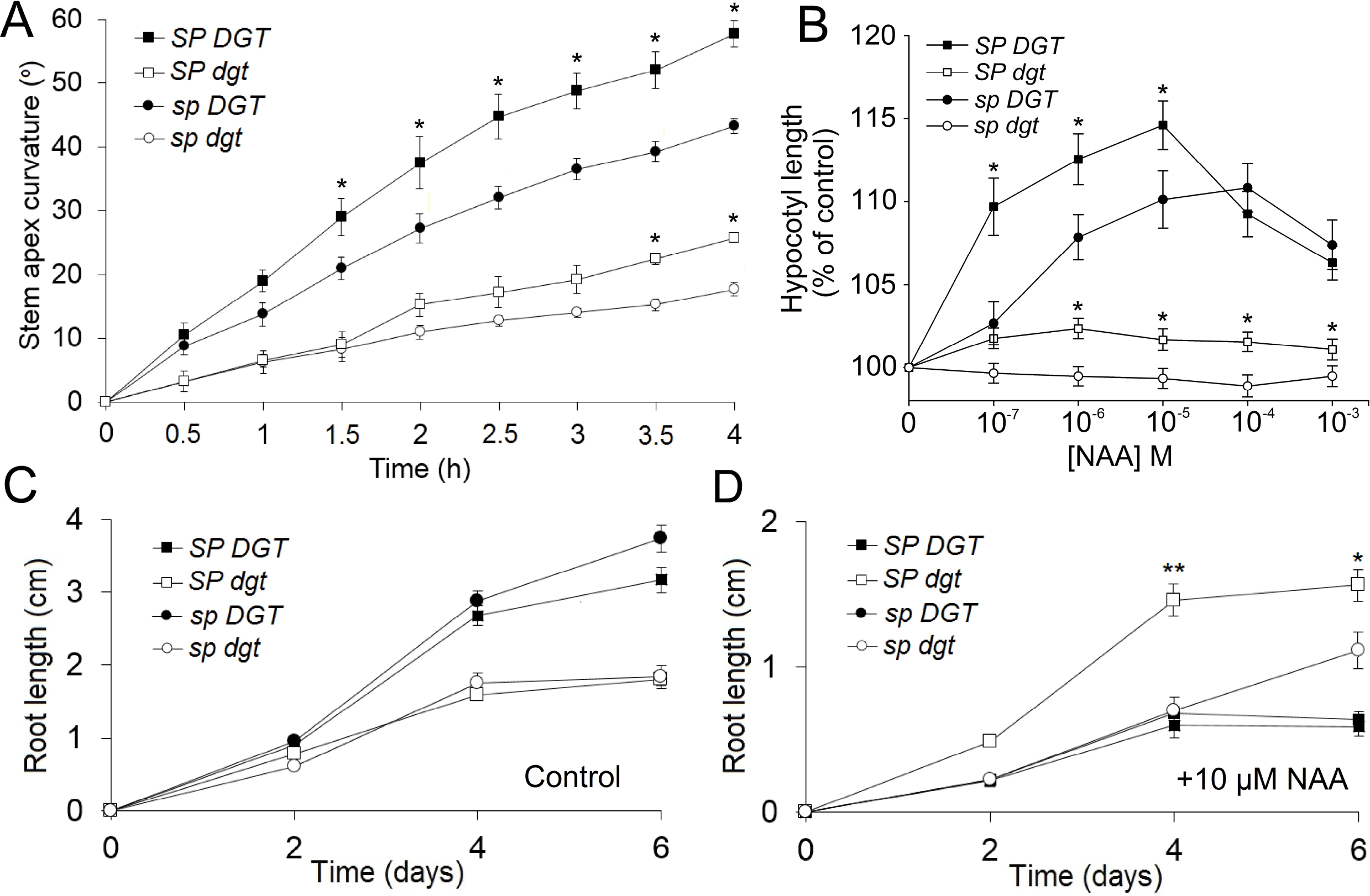
Impact of the *self-pruning* (*sp*)mutation on auxin responses in planta. **(A) Kinetics of gravitropic response in the shoot.** Shoot angle after placing plants horizontally at time point 0 (n=5). (**B) Elongation of excised hypocotyls in response to naphthalenacetic acid (NAA).** 6-mm hypocotyl sections were incubated in the indicated NAA concentration for 24 h before measurement. (n=15) **(C-D) Time-course of *in vitro* root elongation of seedlings in control and 10 μM NAA-containing MS medium.** (n=25). In all panels, bars indicate s.e.m. and asterisks indicate statistically significant differences between *SP* and *sp* plants harboring the same *DIAGEOTROPICA* (*DGT*) allele (*p≤0.05; \*\**p*≤0.01, *t*-test).

**Figure 5.**
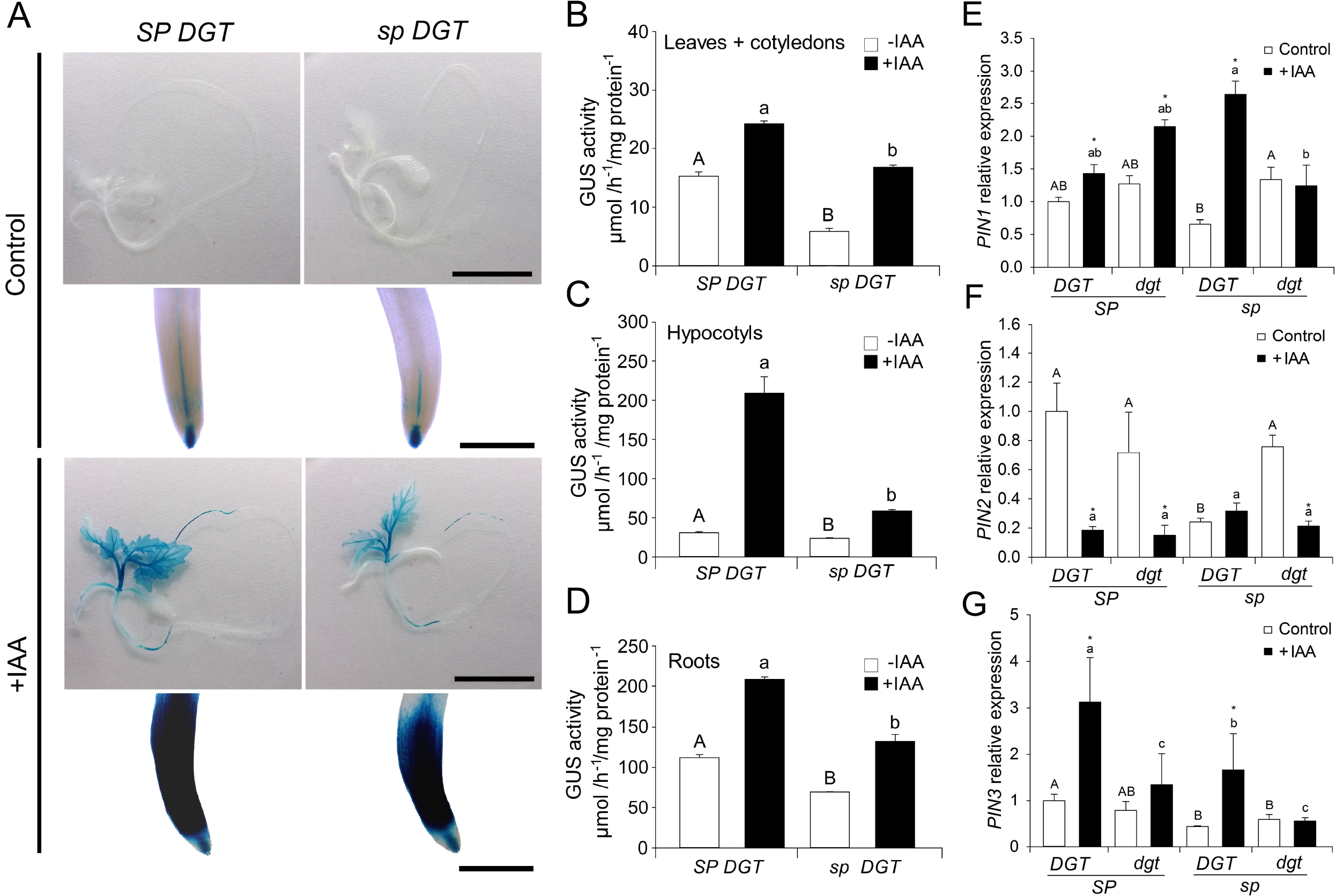
Effects of *SELF-PRUNING* (*SP*) on the auxin signaling and transport machinery *in planta*. **(A) Expression pattern of the GUS reporter driven by the auxin-inducible *DR5* promoter.** Representative wild-type (*SP*) and mutant (*sp*) seedlings (bar=2 cm) and their root tips (bar=250 μm) in the absence or presence of exogenous auxin (20 μM IAA, 3h) 15 dag. **(B-D) Fluorimetric quantification of GUS precipitate.** Seedlings were sampled 15 dag, after treatment with exogenous auxin (20 μM IAA, 3h) or mock. Values are mean ± s.e.m (*n*=4). Letters indicate significant differences between genotypes within the same treatment (p<0.05 ANOVA, Tukey). **(E-G) Relative gene expression of *PIN* transporters in roots.** Letters indicate significant differences between genotypes within the same treatment (p<0.05 ANOVA, Tukey)

Finally, we determined whether auxin affects *SP* at the transcriptional level, as suggested by the presence of auxin-response elements (TGTCTC, and their degenerate version, TGTCNC) (Ulmasov et al., 1995) in the 3′ and 5′ flanking regions of the *SP* gene in tomato and related Solanaceae species (Fig. 6). Analyzing *SP* mRNA levels in seedlings of *SP DGT* and *sp DGT* plants sprayed with IAA or a mock solution, revealed that *SP* expression was induced by IAA treatment in both genotypes. Importantly, *SP* transcript levels were significantly higher in *dgt* mutant plants both in IAA-treated and control seedlings (Fig. 6). We further assessed the effect of *SP* and *DGT* on the mRNA levels of the key players in the auxin signaling cascades, including some members of the *AUXIN RESPONSE FACTORS* (*ARF*) and *AUXIN/INDOLE-3-ACETIC ACID INDUCIBLE* (*Aux/IAA*) gene families, which were chosen by their high auxin-inducibility (Audran-Delalande et al., 2012). *SP* and *DGT* had combinatorial effects in the expression levels of IAA1, *IAA2, IAA9, ARF8*, and *ARF10*, whereas functional *DGT* decreased expression of *IAA3* (Fig. 6).

**Figure 6.**
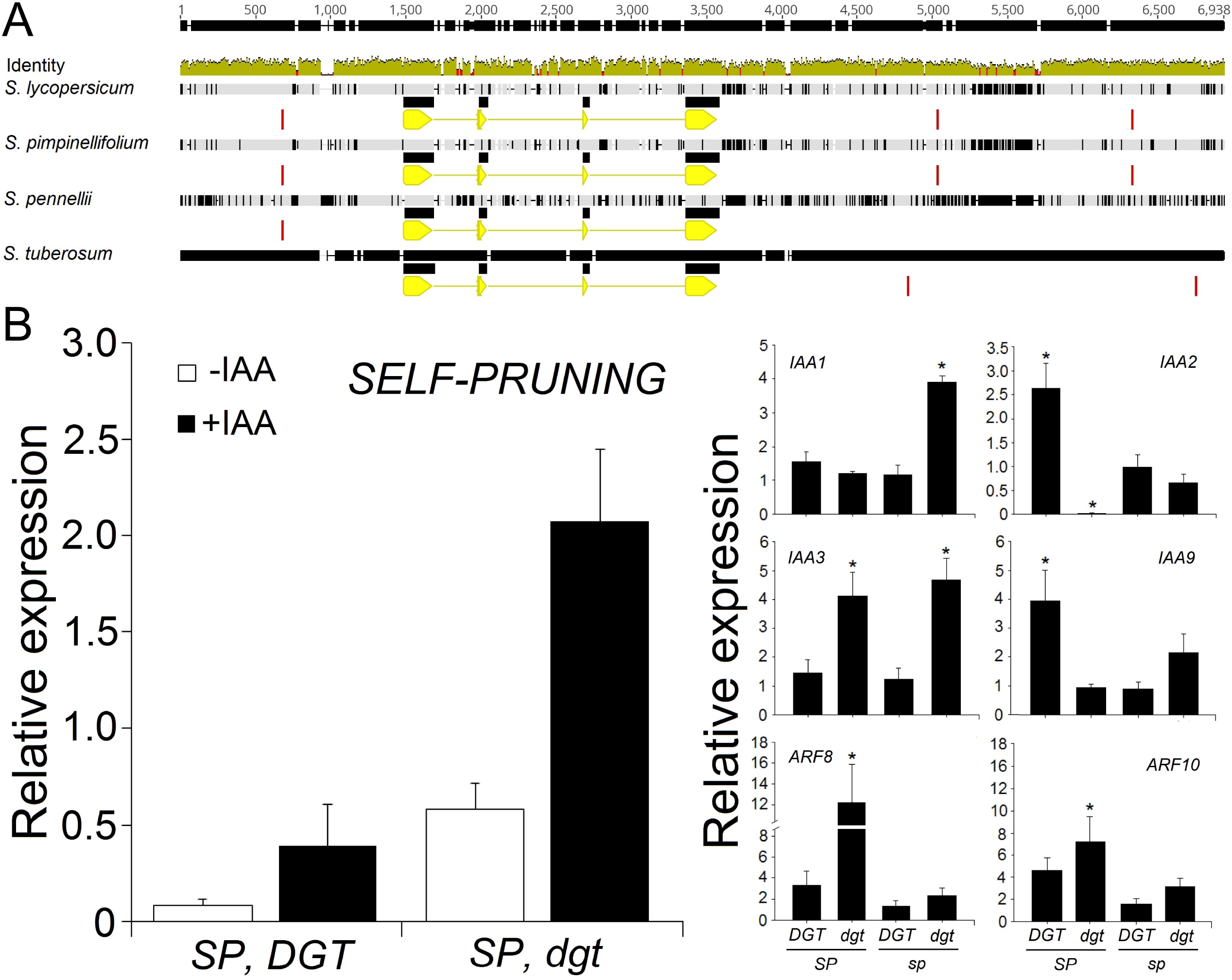
*SELF-PRUNING* (*SP*) and auxin-signaling gene expression is altered by the *diageotropica* (*dgt*) mutation. **(A) Genomic structure of the *SP* gene in solanaceous species:** tomato (*S. lycopersicum*), its wild relatives *S. pimpinellifolium* and *S. pennellii* and potato (*S. tuberosum*). The coding sequence is indicated in yellow (exons, thick bars; introns, thin bars). Red blocks indicate the presence of a conserved or degenerate auxin-response element (AuxRE), TGTCNC. **Relative transcript accumulation of *SP* (B) and auxin signaling genes (C) in sympodial meristems.** Tissues were sampled from 10d old plants, 24h after 10 μM IAA or mock spray. Letters indicate significant differences between genotypes within the same treatment (p<0.05 ANOVA, Tukey). Asterisks indicate significant differences with respect to the wild-type *SP DGT* (p<0.05, t-test).

## Discussion

### Impact of SP alleles, and their interaction with auxin, in the control of shoot architecture

Although it has been previously demonstrated that the *sp* mutation does not alter tomato flowering time or the number of leaves before termination of the shoot (Pnueli et al., 1998; Shalit et al., 2009), *SP* orthologs vary in this respect depending on the species. Flowering time is not affected in *cen* mutants in Antirrhinum (Bradley et al., 1996), whereas Arabidopsis *tfl1* mutants flower earlier and *TFL1* overexpression delays flowering by preventing the meristem transition from vegetative to floral (Ratcliffe et al., 1998). In soybean, where large intraspecific variation exists in time to flowering, association mapping has recently linked this important agronomic trait to the *Dt1* locus, a *CEN/TFL1/SP* ortholog (Zhang et al., 2015). Comparison of determinate and indeterminate near-isogenic soybean lines consistently showed earlier flowering in the former across different locations and planting seasons (Ouattara and Weaver, 1994). Our data show that loss of *SP* function (*sp* allele) leads to slightly but consistently earlier flowering in tomato, measured either in days after germination or the reduction of the number of nodes before the first inflorescence. Using a near isogenic line harbouring the wild type *Dwarf* (D) allele, which codes for a brassinosteroid (BR) biosynthesis gene (Marti et al., 2006), we showed here that this effect is not related to reduced BR levels in MT. This is in agreement with the fact that the phenotypes of the *sp* and *dgt* individual mutants in the MT background closely resemble those published for the same mutations in other tomato cultivars (Carvalho et al., 2011). It is therefore unlikely that the combination of both mutations (*sp* and dgt) would be affected epistatically by the *d* allele (Campos et al., 2010). Interestingly, the *dgt* mutation delays the number of days to flowering in either *SP* or *sp* backgrounds (Balbi and Lomax, 2003), apparently by reducing the expression of *SINGLE FLOWER TRUSS* (*SFT*) gene, which encodes the florigen (Evans, 1971; Shalit et al., 2009). It does not, however, significantly affect the number of leaves produced before termination, which is a proven effect of *SFT* and its orthologs in tomato and other species (Kojima et al., 2002; Lifschitz and Eshed, 2006; Navarro et al., 2015). Hence, the loss-of-function *sft* mutant produced 130% more leaves on the primary shoot than the control MT (Vicente e al., 2015). Conversely, transgenic tomato plants overexpressing *SFT* flower after only three or four leaves (Molinero-Rosales et al, 2004; Shalit et al, 2009).

Axillary branching was increased in *sp* mutants in both *DGT* and *dgt* allele backgrounds. The expression of *SP* is higher in axillary meristems, suggesting a possible role for *SP* in the control of apical dominance (Thouet et al., 2008). Our results reinforce this notion, as *sp* mutants are more profusely branched than wild-type plants. This also agrees with the effects of the *SP* ortholog *Dt1* in soybean, where comparison of determinate and indeterminate isogenic lines revealed an increased propensity to side branching in the former (Gai et al., 1984). The *dgt* mutant responds to auxin treatment of decapitated shoots, which inhibits bud outgrowth to the same extent as in wild-type plants (Cline, 1994). Apical dominance has been reported to be reduced in intact *dgt* plants (Coenen et al., 2003), but caution should be exercised when interpreting these results, as published work on *dgt* has been conducted in tomato cultivars differing in their *SP* alleles (Supplementary Table 1). Our results indicate a strong and complex interaction between *SP* and *DGT* in the control of apical dominance: the *dgt* mutation increased it in the wild-type *SP* background, but also increased axillary bud outgrowth in the *sp* background, enhancing its branching phenotype.

### Control of endogenous auxin levels and polar auxin transport by SP and DGT

Endogenous IAA concentration and distribution within tissues determine a wide range of plant developmental processes, including apical dominance, stem growth, vascular patterning, root development and others (Petrášek and Friml, 2009; Ljung, 2013). IAA synthesis is maximal in younger, developing parts of the plant such as leaflets and root apices (Ljung et al., 2002). IAA levels in dark-grown seedlings (Fujino et al. 1988) and roots (Muday et al., 1995) of *sp dgt* and *SP DGT* plants are indistinguishable. However, free IAA levels in aerial parts of seven-day-old light-grown seedlins of *sp dgt* plants were twice as high as in *SP DGT* plants (Fig. 2), suggesting a light-dependent, synergistic influence of the *sp* and *dgt* alleles on auxin synthesis, degradation, or transport.

The above results could reflect changes in IAA biosynthesis, degradation or transport. The reduction in PAT produced by the *dgt* mutation was described previously (Ivanchenko et al., 2015), but the synergistic effect of the *sp* mutation described here was unexpected. The differences in IAA concentration in the aerial part of the seedlings could be due to altered PAT caused by both the *sp* and *dgt* mutations. PAT from the shoot organs to the root tips induces the formation of the entire plant vascular system (Aloni, 2013; Marcos and Berleth, 2014), as evidenced by the repression of protoxylem formation upon treatment with the auxin transport inhibitor NPA (Bishopp et al, 2012) and the polar localization of PIN1 in pre-procambial cells (Scarpella et al, 2006). Lack of large secondary xylem vessels was conspicuous in the *dgt* mutant, as previously described (Zobel, 1974). In plants harbouring the functional *DGT* allele, the effect of the *sp* mutation was to increase the incidence of larger (>800 μm^2^ cross-sectional area) vessels. In tree species, there is evidence that the relationship between xylem vessel density and size involves differential regulation of the duration of tracheid expansion along the longitudinal (Anfodillo et al, 2012; Sorce et al, 2013) and radial (Tuominen et al, 1997) axes. Tree stature has a strong influence on vessel width due to an allometric scaling effect (Morris et al, 2017). It remains to be seen if this is also the case in herbs, and if the effect of the *SP* gene on xylem width is indirectly caused by its control of plant height, or directly by its influence on PAT. Increased PAT in the *polycotyledon* tomato mutant, for instance, leads to an altered vascular pattern in the hypocotyl (Al-Hammadi et al, 2003; Kharshiing et al, 2010).

### SP affects excised hypocotyl elongation, gravitropic responses, and root regeneration and elongation

Elongation of excised hypocotyl segments in response to different concentrations of exogenous auxin is a classical assay for auxin sensitivity (Gendreau et al., 1997; Collett et al., 2000). The hypocotyl elongation response of *dgt* has been described in the background of tomato cultivar VFN8, a mutant for *sp* (Supplemental Table 1). In both intact or excised hypocotyl segments, a reduced response to exogenous auxin was observed for the *sp dgt* double mutant (Kelly and Bradford, 1986; Rice and Lomax, 2000). We confirmed these results, but show that a functional *SP* allele leads to increased elongation in either *DGT* or *dgt* backgrounds. Collectively, these results indicate that some compensatory effect can be ascribed to SP on this response. Hypocotyl elongation in Arabidopsis relies on auxin-induced changes in the activity of plasma membrane H^+^-ATPases, which leads to increased H^+^ extrusion and cell expansion, through expansin-mediated cell wall loosening, as per the acid growth theory (Takahashi et al., 2012). Hypocotyl elongation upon exogenous auxin application, points to a positive effect of SP on the activity of plasma membrane H^+^-ATPases. Interestingly, the activity of both plasma membrane H^+^-ATPases and PIN efflux transporters, which are also influenced by SP at the transcriptional level (Fig. 4), is regulated by changes in their phosphorylation state (Takahashi et al., 2012; Zourelidou et al., 2014; Weller et al., 2017). This fits with earlier suggestions that *SP*, which encodes a phosphatidylethanolamine binding protein (PEBP), exerts at least some of its effects on membrane proteins through interaction with kinases (Pnueli et al., 2001).

The Cholodny-Went hypothesis is a classical model suggesting that differential auxin distribution is the cause of directional plant bending with respect to an exogenous stimulus such as light or gravity (Went, 1974). That DGT is required for a correct gravitropic response of roots and shoots has been demonstrated, but the explanation at the molecular level is still lacking (Muday et al., 1995; Rice and Lomax, 2000). To the best of our knowledge, an effect of SP on shoot gravitropism had not been tested before. Functional SP enhances shoot gravitropism in horizontally positioned plants of either *DGT* or *dgt* background. Functional SP produces taller plants, so it is tempting to speculate that they should have a stronger gravitropic response in order to facilitate the bending of a larger stem. In Arabidopsis, the IAA efflux transporter PIN3 mediates lateral redistribution of auxin and is therefore involved in hypocotyl and root tropisms (Friml et al., 2002). It seems reasonable to suggest a link between SP and PIN3 in the face of our *PIN* gene expression profiles (Fig. 5). Remarkably, both types of efflux transporters, PIN1 and PIN3, have been shown to relocate at the subcellular level via the same mechanism: vesicle trafficking along the actin cytoskeleton between the plasma membrane and endosomes (Geldner et al., 2001; Friml et al., 2002). Dissecting the intertwined mechanisms involved in this possible co-regulation will be required to fully understand to which extent and how exactly SP affects auxin distribution.

High concentrations of exogenous auxin inhibit root elongation. As expected, the *dgt* mutation reduced auxin-induced inhibition (Coenen and Lomax, 1998) in root elongation, however, a functional *SP* allele led to lower inhibition than in the double *sp dgt* mutant. This result could be ascribed to a new balance in auxin transport and signaling produced by the combination of *SP* and *DGT*. Root growth increases with the strength of auxin signaling up to a certain optimum, and then begins to decline, probably following a parabolic trajectory (Sibout et al, 2006). *In vitro* root regeneration, on the other hand, is stimulated by low concentrations of auxin and *dgt* is relatively insensitive to this exogenous treatment (Coenen and Lomax, 1998). Interestingly, the *sp* mutation also reduces rhizogenesis (Fig S3), which reinforces the notion of *SP* positively influencing PAT, as the PAT inhibitor TIBA has been shown to reduce *in vitro* root formation in tomato (Tyburski and Tretyn, 2004).

### Interactions between SP and the auxin signaling machinery

Auxin signaling output can be estimated by following *DR5* promoter activation pattern (Ulmasov et al., 1995; Liao et al., 2015). For example, GUS staining of *DR5::GUS* has revealed that auxin flux at the root tips proceeds acropetally up to the root cap, where it is redistributed via lateral efflux transporters toward a peripheral basipetal transport route (Benková et al., 2003; Paciorek et al., 2005; Dhonukshe et al., 2008). Exogenous auxin application leads to greater GUS signal in seedlings with a functional *SP* allele, owing probably to alterations in the auxin signaling machinery produced by SP, such as expression of *Aux/IAA* and *ARF* family genes (Fig. 6).

Auxin signalling is strongly dependent on auxin levels and the responsiveness of target cells. At low IAA levels, a suite of repressor proteins, including Aux/IAA and TOPLESS, repress ARFs, a group of transcription factors which regulate the expression of auxin-responsive genes (Causier et al., 2012; Bargmann and Estelle, 2014; Chandler, 2016). At high IAA levels, auxin acts as a molecular glue to stabilize the TIR1/AFB receptor binding and tagging of Aux/IAAs for 26S proteasome degradation (Hayashi, 2012). This, in turn, frees ARFs bound to Auxin-Response Elements (AuxRE) in the genome (TGTCTC, or its degenerate, but also functional form, TGTCNC) to activate or repress gene expression (Ulmasov et al., 1995). Our *in silico* analyses demonstrated the presence of conserved AuxRE elements both 5′ (upstream) and 3′ (downstream) of the *SP* coding sequence in the genome of tomato and closely related species. It has recently been shown that the 3′ region of *TFL1* in Arabidopsis contains multiple auxin *cis*-regulatory elements key for the control of spatio-temporal expression of the gene (Serrano-Mislata et al., 2016). It remains to be determined whether such *cis*-regulatory elements are also functional in tomato and if they are involved in the response of *SP* expression to auxin.

Tomato has 25 *Aux/IAA* and 22 *ARF* genes (Audran-Delalande et al., 2012; Zouine et al., 2014), indicating that auxin signalling is very complex. DGT can alter the expression of genes related to auxin signaling (Mito and Bennett, 1995), including *Aux/IAA* genes (Nebenführ et al., 2000). It was recently discovered that cyclophilin peptidyl-prolyl isomerases (PPIases) catalyse the *cis/trans* isomerisation of peptide bonds preceding proline residues of target peptides, including Aux/IAAs (Jing et al., 2015). Only Aux/IAA peptides of the right conformation can bind to the TIR1 receptor and be tagged for degradation, thus PPIases, such as DGT, are believed to play a key role in auxin perception (Su et al., 2015). It is likely that some transcriptional feedback exists when the right conformers are not produced, as suggested by increased transcript levels of *IAA3* and reduced levels of *IAA9* in *dgt* mutants. Furthermore, our *in silico* analysis shows that the *sp* mutation occurs in a highly conserved cis-proline residue in a DPDxPxn10H consensus region in the PEBP domain (Fig. S3), which is a potential target for PPIases. Whether this putative molecular interaction between SP and DGT could account for the phenotypic outcomes shown here, remains to be determined.

## Conclusions

Auxin gradients are critical for organogenesis in the shoot apex, however, the influence of this hormone on shoot determinacy, which is a key determinant of growth habit, has never been addressed in depth. Our data provides the first link between auxin and the anti-florigenic protein SELF-PRUNING (SP), the main switch between indeterminate and determinate growth habit in tomato. Although it is not clear whether auxin itself can affect growth habit, a physiological interaction between this hormone and members of the CETS family was clearly demonstrated here. Hence, *SP* alleles affected various auxin-related responses (*e.g*. apical dominance, PIN1-mediated polar auxin transport, vascular differentiation, H^+^ extrusion and gravitropism responses), different *SP* orthologs presented AuxREs, and the auxin mutant *dgt* downregulated *SFT* and upregulated *SP* expression. There are now increasing evidences that *SP/SFT* genetic module is a hub in crop productivity, affecting heterosis for yield (Krieger *et al.*, 2010) and improving plant architecture and the vegetative-to-reproductive balance (McGarry and Ayre, 2012; Vicente et al., 2015; Zsögön *et al.*, 2017). Our results suggest that at least part of the effect of the SP/SFT module on yield is mediated by auxin. This knowledge may inspire novel and more precise manipulation of this hormone for applications in agriculture.

## Materials and methods

### Plant material

Seeds of the tomato (*Solanum lycopersicum*, L.) cultivar Micro-Tom (MT) were kindly donated by Dr. Avram Levy (Weizmann Institute of Science, Israel) in 1998 and subsequently maintained (through self-pollination) as a true-to-type cultivar. MT seeds carrying the synthetic auxin-responsive (*DR5*) promoter fused to the reporter gene *uid* (encoding a β-glucuronidase, GUS) were obtained from Dr. José Luiz García-Martínez (Universidad Politécnica de Valencia, Spain). The *diageotropica* (*dgt*) mutation was introgressed into MT from its original background in cv. VFN8 (LA1529), donated by Dr. Roger Chetelat (Tomato Genetics Resource Center, Davis, University of California, USA). The functional allele of *SELF-PRUNING* was introgressed from cv. Moneymaker (LA2706).

Introgression of mutations into the MT cultivar was described previously (Carvalho et al., 2011). A comparison between indeterminate (*SP/SP*) and determinate (*sp/sp*) plants in the MT background has been published previously (Vicente et al., 2015). Both *sp* and *dgt* mutations were confirmed by CAPS marker analyses and sequencing, respectively. All experiments were conducted on BC_6_F_3_ plants or subsequent generations (Sestari et al., 2014). *In vitro* seedling cultivation was conducted under controlled conditions (16h/8 h day/night, approximately 45 μmol m^−2^ s^−1^ PAR, 25±1^0^C) in flasks with 30 ml of MS/2 media gellified with 0.5% agar, pH 5.8. Seeds were surface sterilized by agitation in 30% (v/v) commercial bleach (2.7% sodium hypochlorite) for 15 min followed by three rinses with sterile distilled water.

### Growth conditions

Plants were grown in greenhouse in Viçosa (642 m asl, 20°45’ S; 42°51’ W), Minas Gerais, Brazil, under semi-controlled conditions: mean temperature of 28°C, 11.5 h/13 h (winter/summer) photoperiod, and 250-350 μmol m^−2^ s^−1^ PAR irradiance and irrigation to field capacity twice a day. Seeds were germinated in 350-mL pots with a 1:1 (v/v) mixture of commercial potting mix Basaplant® (Base Agro, Brazil) and expanded vermiculite supplemented with 1g L^−1^ 10:10:10 NPK and 4 g L^−1^ dolomite limestone (MgCO_3_ + CaCO_3_). Upon appearance of the first true leaf, seedlings of each genotype were transplanted to pots containing the soil mix described above, except for the NPK supplementation, which was increased to 8 g L^−1^.

### Auxin quantification

Endogenous indole acetic acid (IAA) levels were determined by gas chromatography tandem mass spectrometry-selecting ion monitoring (GC-MS-SIM, Shimadzu model GCMS-QP2010 SE). Samples (50-100 mg fresh weight, FW) were extracted and methylated as described in (Rigui et al., 2015). About 0.25 μg of the labeled standard [^13^C_6_]IAA (Cambridge Isotopes, Inc.) was added to each sample as internal standards. The chromatograph was equipped with a fused-silica capillary column (30m, ID 0.25 mm, 0.50 μm thick internal film) DB-5 MS stationary phase using helium as the carrier gas at a flow rate of 4.5 mL min^−1^ in the following program: 2 min at 100°C, followed a ramp by 10°C min^−1^ to 140°C, 25°C min^−1^ to 160°C, 35°C min^−1^ to 250°C, 20°C min^−1^ to 270°C and 30°C min^−1^ to 300°C. The injector temperature was 250°C and the following MS operating parameters were used: ionization voltage, 70 eV (electron impact ionization); ion source temperature, 230°C; interface temperature, 260°C. Ions with a mass ratio/charge (m/z) of 130 and 189 (corresponding to endogenous IAA) and 136 and 195 (corresponding to [^13^C_6_]-IAA) were monitored and endogenous IAA concentrations were calculated based on extracted chromatograms at m/z 130 and 136.

### Polar auxin transport analysis

Polar auxin transport (PAT) was assayed in hypocotyl segments of 2-week-old seedlings according to the protocol originally described by (Al-Hammadi et al., 2003) with some modifications. Briefly, 10 mm hypocotyl sections were excised and incubated in 5 mM phosphate buffer (pH 5.8) containing 1 μM IAA for 2 h at 25 ± 2°C on a rotary shaker (200 rpm). These segments were placed between receiver (1% [w/v] agar in water) and donor blocks (1% [w/v] agar in 5 mM phosphate buffer [pH 5.8] containing 1 μM IAA and 100 nM ^3^H-IAA) oriented with their apical ends toward the donor blocks. After 4 h of incubation inside a humid chamber at 25 ± 2°C, the receiver blocks were removed and stored in 3 mL scintillation cocktail (Ultima Gold™, PerkinElmer, USA). Receiver blocks plus scintillation cocktail were shaken overnight at 100 rpm and 28 ± 2°C before analysis in a scintillation counter. As negative control, some hypocotyl segments were sandwiched for 30 min between 1-*N*-natphthylphthalamic acid (NPA)-containing blocks (1% [w/v] agar in water containing 20 μM NPA) prior the auxin transport assays. ^3^H d.p.m. was converted to fmol auxin transported as described in (Lewis and Muday, 2009).

### Auxin sensitivity assays

Root regeneration from cotyledon explants was conducted as described (Cary et al, 2001). Briefly, cotyledon explants were obtained from eight-day old seedlings germinated *in vitro* in half-strength MS medium. The explants were then incubated on Petri dishes containing MS with or without supplementation with 0.4 μM NAA. After 8 days, the number of explants with visible roots (determined using a magnifying glass) was counted.

For hypocotyl elongation assays, hypocotyls were excised from two week-old seedlings and cut into 5-mm sections. Between 15 and 20 segments were pre-incubated for each treatment, floated on buffer (10 mM KCl, 1 mM MES-KOH [pH 6.0], and 1%[w/v] Suc for 2 h at 25°C in the dark to deplete endogenous auxin. Segments were then incubated on buffer (10 mM KCl, 1 mM MES-KOH [pH 6.0], and 1%[w/v] Suc and NAA at the indicated concentration for 24 h on a shaker at 25°C under white light. Segments were photographed to determine their length using ImageJ (NIH, Bethesda, MA, USA). The experiment was repeated three times with similar results.

For the gravitropism assays, plants were germinated in 350 ml pots and transferred to 50 mL Falcon tubes two days afer germination (dag). The gravitropic response was assessed 10 dag by placing five plants of each genotype horizontally and photographing them in 30 minute intervals. The angle of shoot bending at each time point was determined using AutoCad 2016 (Autodesk, Inc., San Rafael, CA, Estados Unidos da Amèrica). Sterilized seeds were germinated in petri dishes onto two layers of filter paper moistened with distilled water, and incubated for 4 d at 25°C in the dark. Ten germinated seeds with radicles of 5-10 mm were transferred to vertically oriented square Petri dishes (120 mm × 120 mm) aligned on each plate with the radicles pointing down. The plates contained MS medium supplemented with vitamins, pH 5.7, 3% (w/v) sucrose, 0.8% (w/v) agar and 10 μM α-naphthaleneacetic (NAA) for the auxin treatment. Plates were incubated in a growth chamber in the dark.

*In vitro* root elongation in response to exogenous auxin was assessed as follows. Seeds were surface-sterilized and imbibed for two days at 4°C in the dark on agar plates containing half-strength MS growth medium (Murashige and Skoog, 1962), transferred to growth chamber under control conditions (12h photoperiod, 150 μmol m^−2^ s^−1^ white light, 22°C/20°C throughout the day/night cycle, 60% relative humidity). After four days, ten seedlings per plate were transferred to half-strength MS medium with or without 10μM NAA (α - naphthalene acetic acid - Sigma-Aldrich, St. Louis, MO, USA) and covered completely with aluminum foil for eight days. Root elongation was assessed every second day under dim light conditions.

### Histochemical assays

Transgenic *DR5::GUS* plants were incubated overnight at 37°C in GUS staining solution (100 mM NaH_2_PO_4_; 10 mM EDTA, 0,5 mM K_4_Fe(CN)_6_; 0,05% Triton X-100, 1mM 5-bromo-4-chloro-3-indolyl-beta-D-glucuronic acid). Following GUS staining, samples were washed in a graded ethanol series to remove chlorophyll. Samples were then photographed using a Leica S8AP0 (Wetzlar, Germany) magnifying glass set to 80× magnification, coupled to a Leica DFC295 camera (Wetzlar, Germany). Quantitative GUS activity was assayed according to Jefferson et al. 1987, with some modifications. Briefly, samples were ground in liquid nitrogen and subsequently homogenized in MUG extraction buffer composed of 50 mM Hepes-KOH (pH 7.0), 5 mM DTT and 0.5% (w/v) PVP. After centrifugation, 200 μL aliquots of the supernatant was mixed with 200 μL GUS assay buffer composed of 50 mM HEPES-KOH (pH 7.0), 5 mM DTT, 10 mM EDTA and 2 mM 4-methylumbelliferyl-β-D-glucuronide (MUG) and incubated at 37 °C for 30 minutes. Subsequently, aliquots of 100 μL were taken from each tube and the reactions were stopped and fluorescence was analyzed using a spectrofluorometer (LS55, Perkin Elmer) with 365 nm excitation and 460 nm emission wavelength (5 nm bandwidth).

### Gene expression analyses

Total RNA was extracted from approximately 30 mg FW of sympodial meristems of 10-day old plants following the protocol of the manufacturer (Promega SV total RNA isolation sytem). For auxin treatments plants were previously sprayed with 10 μM IAA or mock-sprayed 24 h prior to RNA extraction. Four biological replicates were used for subsequent cDNA synthesis. Each replicate consisted of a pool of three plants each were used for the analyses, since sympodial meristems are small and did not provide enough biological material for RNA extraction. Two technical replicates were then performed on each of the four samples. RNA integrity was analyzed on 1% agarose gel and RNA concentration was estimated before and after treatment with DNase I (Amplification Grade DNase I, Invitrogen). Total RNA was transcribed into cDNA using the enzyme reverse transcriptases, Universal RiboClone® cDNA Synthesis (Promega, Madison, WI, USA) following the manufacturers’ protocols.

For gene expression analyses Power SYBR® green PCR Master Mix was used in MicroAmp™ Optical 96-well reaction plates (both from Applied Biosystems, Singapore) and adhesive film MicroAmp™ Optical (Applied Biosystems, Foster City, CA, USA). The number of reactions from the cycle threshold (CT) as well as the efficiency of reaction were estimated using the Real-Time PCR Miner tool (Zhao and Fernald, 2005).

Relative expression was normalized using actin and ubiquitin; actin was used to calculate ΔΔCT assuming 100% efficiency of amplification of genes (2^- ΔΔCT). Primer sequences used are shown in Supplementary Table 1. Melting curves were checked for unspecific amplifications and primer dimerization.

### *In silico* sequence analyses

*SP* gene alignments was performed using the ClustalW alignment option of the Geneious R9 (Biomatters, Auckland, New Zealand) software package.

### Statistical analysis

ANOVA and Tukey HSD tests were performed using Assistat 7.6 beta (http://assistat.com). Percentage data were converted to inverse function (1/X) before analysis.

## Acknowledgments

This work was supported by funding from the Agency for the Support and Evaluation of Graduate Education (CAPES-Brazil), the National Council for Scientific and Technological Development (CNPq-Brazil), Foundation for Research Assistance of the São Paulo State (FAPESP-Brazil) and the Foundation for Research Assistance of the Minas Gerais State (FAPEMIG-Brazil). We thank CAPES for studentships granted to J.M.R., W.B.S., and D.S.R. FAPESP provided grants for M.H.V. (2016/05566-0), L. F. (2013/18056-2), L.E.P.P. (2015/50220-2), and A.Z. (2013/11541-2). W.L.A. and L.E.P.P. also acknowledge grants from CNPq (grant 307040/2014-3 to L.E.P.P.). We thank Biomatters Ltd. (Auckland, New Zealand) for the kind gift of a Geneious R9 licence.

**Author contributions**
WBS, MHV and JMR generated the plant material and conducted experiments. WBS, MHV, JMR, DSR, LF, RCF and RB conducted experiments and prepared figures and/or tables. LF and WLA designed experiments, contributed reagents/materials/analysis tools and reviewed drafts of the paper. AZ and LEPP conceived and designed the experiments, analyzed the data and wrote the paper.

